# Dynamic Erasure of Random X-Chromosome Inactivation during iPSC Reprogramming

**DOI:** 10.1101/545558

**Authors:** Adrian Janiszewski, Irene Talon, Juan Song, Natalie De Geest, San Kit To, Greet Bervoets, Jean-Christophe Marine, Florian Rambow, Vincent Pasque

**Affiliations:** KU Leuven - University of Leuven, Department of Development and Regeneration, Herestraat 49, B-3000 Leuven, Belgium.; Laboratory for Molecular Cancer Biology, VIB Center for Cancer Biology; Department of Oncology, KU Leuven, Belgium

**Keywords:** Epigenetic reprogramming, X-chromosome reactivation, Epigenetic memory, iPSCs, Chromatin silencing

## Abstract

**Background:** Induction and reversal of chromatin silencing is critical for successful development, tissue homeostasis and the derivation of induced pluripotent stem cells (iPSCs). X-chromosome inactivation (XCI) and reactivation (XCR) in female cells represent chromosome-wide transitions between active and inactive chromatin states. While XCI has long been studied and provided important insights into gene regulation, the dynamics and mechanisms underlying the reversal of stable chromatin silencing of X-linked genes are much less understood. Here, we use allele-specific transcriptomic approaches to study XCR during mouse iPSC reprogramming in order to elucidate the timing and mechanisms of chromosome-wide reversal of gene silencing.

**Results:** We show that XCR is hierarchical, with subsets of genes reactivating early, late and very late. Early genes are activated before the onset of late pluripotency genes activation and the complete silencing of the long non-coding RNA (lncRNA) *Xist*. These genes are located genomically closer to genes that escape XCI, unlike those reactivating late. Interestingly, early genes also show increased pluripotency transcription factor (TF) binding. We also reveal that histone deacetylases (HDACs) restrict XCR in reprogramming intermediates and that the severe hypoacetylation state of the Xi persists until late reprogramming stages.

**Conclusions:** Altogether, these results reveal the timing of transcriptional activation of mono-allelically repressed genes during iPSC reprogramming, and suggest that allelic activation involves the combined action of chromatin topology, pluripotency transcription factors and chromatin regulators. These findings are important for our understanding of gene silencing, maintenance of cell identity, reprogramming and disease.

## INTRODUCTION

Development, tissue homeostasis and the derivation of iPSCs depend on the accurate establishment, maintenance and reversal of chromatin silencing. While the formation of facultative heterochromatin has been extensively studied (1), it remains unclear how epigenetic memory of stable gene silencing is reversed by TFs and accompanying chromatin mechanisms. Chromosome-wide transitions between active and inactive chromatin states are excellently modeled by XCI and XCR in female mammals (2-7). XCI involves epigenetic mechanisms including lncRNAs, chromatin modifications and changes in chromosome topology and leads to the establishment and maintenance of stable gene silencing (1, 8, 9). However, little is known about how cells erase epigenetic memory of stable silenced chromatin.

XCI is the rapid, chromosome-wide silencing of an entire X-chromosome during female mammalian development. It ensures dosage compensation between XX female and XY male cells (10). Moreover, XCI involves allelic gene regulation resulting in mono-allelic expression, a phenomenon shared with several genes on autosomes, such as imprinted genes (11). In early mouse embryos, imprinted XCI (iXCI), which always inactivates the paternal X-chromosome, takes place at 4-cell stage. It is followed by XCR, in the inner cell mass (ICM) of the blastocyst (12-15). Then, random XCI (rXCI) of one of the two X-chromosomes is induced, to establish dosage compensation in the epiblast (10).

The XCI process is initiated by the lncRNA *Xist* and leads to the removal of active histone marks such as histone acetylation and the accumulation of repressive histone marks such as histone H3 lysine 27 trimethylation (H3K27me3) on the future Xi (16-18, 19l, 20). Moreover, recent studies on neural progenitor cells show that, upon inactivation, the Xi undergoes conformational changes which include both loss of topologically associated domains (TADs) and the subsequent folding into two silenced mega-domains (8). Remarkably, a subset of X-linked genes escapes XCI and maintains bi-allelic expression. These genes, termed “escapees”, bypass the suppressive effects of *Xist* and the repressive protein complexes, and are located within the residual TAD-like clusters on the Xi (8, 21). XCI, therefore, provides a paradigmatic example of chromosome-wide gene silencing stably maintained in somatic cells. The precise relationship between epigenetic modifications on the randomly inactivated X-chromosome and the stability and reversibility of gene silencing remains unclear.

The epigenetic memory of the Xi can be reversed by the process of XCR. During development, XCR takes place in the epiblast in mouse and during the formation of female primordial germ cells in mouse and human (12, 13, 22). Despite its importance, much less is known about XCR compared to XCI. Opposite to XCI, XCR leads to the silencing of *Xist*, increased expression of antisense lncRNA *Tsix*, loss of repressive chromatin marks, recruitment of active chromatin modifications and, eventually, chromosome-wide gene reactivation (23). Previous studies during mouse development have shown that the reversal of iXCI is a rapid but gradual process that initiates before the loss of *Xist* and is partly restricted by H3K27me3 which is actively removed by UTX H3K27 histone demethylase in order to activate slowly reactivating genes (15). However, the dynamics and mechanisms that mediate reversal of rXCI, as opposed to the reversal of iXCI, remain to be elucidated.

Reprogramming of female somatic cells into iPSCs induces chromosome-wide erasure of gene silencing from rXCI. Previous studies on reprogramming to iPSCs have shown that XCR occurs late (24), after the silencing of *Xist* and the activation of pluripotency markers such as NANOG and DPPA4 (25-27). *Xist* deletion did not affect these kinetics, however, its ectopic expression caused a delay in XCR suggesting that silencing of *Xist* is required for XCR (26). Additionally, inducing pluripotency by cell fusion of human female fibroblasts with mouse embryonic stem cells (ESCs) leads to partial XCR (28). These studies suggested that there might be different levels of susceptibility of silenced X-linked genes to reactivation (28, 29). Nevertheless, it is still unknown whether the kinetics of reactivation during reprogramming to iPSCs vary for different genes and what are the factors and chromatin features that enable or restrict XCR.

Pluripotency has been strongly linked to XCR. The presence of two active X-chromosomes is considered a conserved hallmark of naïve pluripotency from mice to humans (30, 31). Previous studies have established a clear link between the presence of a robust pluripotency network and stable suppression of *Xist* expression (32-35). However, the mechanisms linking pluripotency factors to *Xist* repression, and XCR, have been unclear. One study proposed that pluripotency factors repress *Xist* by direct binding to *Xist* intron 1 (32). However, deletion of this region had little effects on XCI or XCR (36). As a result, it is still not known how pluripotency factors mediate XCR and bi-allelic X-linked gene expression. Given the importance of TFs in transcriptional regulation, one hypothesis is that pluripotency TFs may, in addition to repressing *Xist* via unknown regulatory regions, also directly bind X-linked genes for their transcriptional activation in the pluripotent state, but until now no evidence to lend support for such a model has been reported.

The reversibility of gene silencing might also depend on the architecture of the Xi. Escapee genes have been shown to be located outside of the two repressive mega-domains of the Xi which allows their bi-allelic activity (8). This raises the possibility that the location of repressed genes in 3D space might be linked to the stability of gene silencing. Yet, it remains unknown whether the location of genes on the X-chromosome, relative to suppressed compartments, is relevant for the reversal dynamics of rXCI.

In this study, we define the kinetics of chromosome-wide X-linked gene reactivation during reprogramming of mouse somatic cells into iPSCs. We ask which genomic and epigenetic marks correlate with the timing of X-linked gene reactivation. In addition, we aim to identify the mechanisms that may enable or restrict reversal of gene silencing during cell fate conversion. We also test how functional interference with chromatin pathways influences XCR upon entry into pluripotency. Our study identifies gene regulatory principles that may ensure the stability of repressed chromatin, potentially applicable in other contexts. Such principles may also facilitate stable maintenance of cellular identity. Finally, this study provides a framework for how TFs induce reversal of stable gene silencing by overcoming active chromatin barriers in order to activate transcription and reverse epigenetic memory.

## RESULTS

### Allele-specific transcriptional analyses during somatic cell reprogramming to induced pluripotency

Overexpression of TFs *Oct4*, *Sox2*, *Klf4* and *c-Myc* (OSKM) in somatic cells leads to the induction of pluripotency and the erasure of transcriptional silencing of the Xi (24). However, the precise timing and underlying mechanisms of chromosome-wide transcriptional activation of X-linked genes during the reversal of rXCI remain to be determined. To define the transcriptional dynamics of XCR, we established an inducible reprogramming mouse model suitable for allele-resolution transcriptome studies. We first isolated female mouse embryonic fibroblasts (MEFs) from highly polymorphic mouse strains originating from the cross between female *Mus musculus musculus* (*Mus*) with a X-GFP transgene on the X-chromosome (37) and male *Mus musculus castaneus* (*Cast*) mice, carrying high density of single nucleotide polymorphisms (SNPs) spread throughout the genome (Figure 1A). To ensure that the starting cells carry the same Xi, we used fluorescence activated cell sorting (FACS) to specifically select only the female MEF cells with silenced X-GFP allele (GFP-negative cells) (Figure 1A, S1A, B). We then induced the reprogramming of Xi-GFP female MEFs into iPSCs by overexpression of (OSKM). Reprogramming led to the appearance of iPSC colonies that then reactivated the Xi as judged by time-resolved live imaging of GFP activation (Figure 1B, S1C). Thus, our strategy enabled conditional reprogramming of somatic cells into iPSCs, accompanied by XCR, using a system compatible with allele-resolution genomic studies.

**Figure 1.**
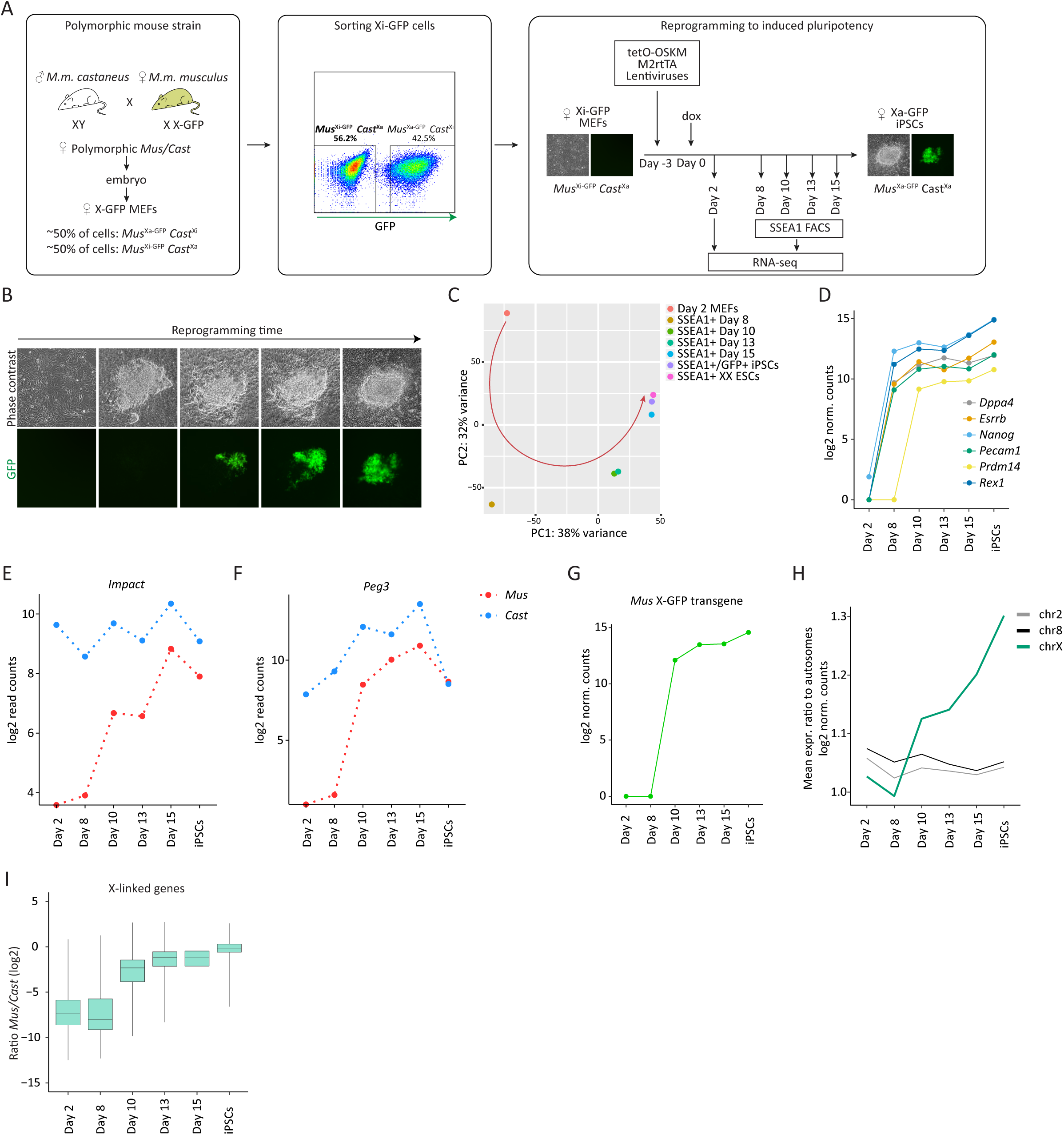
Allele-resolution system to study transcriptional reactivation of X-linked genes during reprogramming to pluripotency. (A) Schematic representation of the system used to trace XCR during reprogramming. Female mice from *M.m. musculus* with a GFP transgene on the X-chromosome were crossed with male mice from *M.m. castaneus.* Next, *Mus/Cast* polymorphic female MEFs were isolated from the offspring, with maternally derived X-GFP by sorting for GFP-(*Mus*^Xi-GFP^/*Cast*^Xa^) cells. Doxycycline (dox) was used to induce the expression of the conditional cassette with OSKM (tetO-OSKM). M2rtTA = reverse tetracycline transactivator. (B) Phase contrast and fluorescent images of representative stages of reprogramming from day 0 to day 15, starting from female *Mus*^Xi-GFP^/*Cast*^Xa^ MEFs to *Mus*^Xa-GFP^/*Cast*^Xa^ iPS cells. GFP+ cells are increasing in number as a result of XCR. (C) Principal component analysis (PCA) of gene expression from different stages of reprogramming. Each colored dot represents a different time point: day 2 (orange), 8 (dark yellow), 10 (green), 13 (turquoise), 15 (blue), SSEA1+/GFP+ iPSCs after four passages (purple) and female ESCs (pink). (D) Levels of pluripotency network gene expression (log2 transformed normalized read counts) during the time course of reprogramming. (E) Expression levels of *Impact*, an autosomal, paternally imprinted gene during factor-induced reprogramming. (F) As in (E) for *Peg3*. (G) X-GFP transgene expression during reprogramming (log2 transformed normalized read counts). (H) Mean expression ratio of chromosome 2, 8 and X relative to autosomes (log2 normalized counts) during reprogramming. (I) X-linked genes expression ratio *Mus/Cast* (log2 transformed normalized read counts). See methods.

To define the kinetics of XCR with allele-resolution, we isolated, at different time points, reprogramming intermediates marked by reactivation of the cell surface marker SSEA1 (Figure S1C). SSEA1 has been shown to mark cells poised for successful acquisition of the pluripotency program and XCR (38-40). The first SSEA1 positive (+) cells isolated did not show significant GFP fluorescence, then gradually reactivated GFP, while fully reprogrammed iPSCs were mostly GFP+ (Figure S1C). We applied full transcript RNA-seq Smart-seq2 to populations of day 2 MEFs, SSEA1+ reprogramming intermediates obtained at day 8, day 10, day 13 and day 15 of reprogramming, as well as iPSCs and control ESCs. We observed gradual changes in the transcriptome of SSEA1+ intermediates, and found that, transcriptionally, day 15 intermediates closely resembled iPSCs and ESCs, indicating successful reprogramming (Figure 1C).

Reprogramming to iPSCs and entrance into the pluripotency program are concomitant with the activation of key pluripotency genes (41). Indeed, pluripotency markers such as *Nanog, Esrrb, Rex1, Pecam1* and *Dppa4* were expressed in day 8 SSEA1+ cells (Figure 1D). However, *Prdm14* was activated only later, at day 10 (Figure 1D). Next, we set out to verify whether this strategy allows retaining allele-specific information and monitored allelic expression of imprinted genes during reprogramming. We examined the expression of *Impact* and *Peg3.* As expected, *Impact* and *Peg3* were expressed exclusively from paternal alleles in MEFs (42, 43), validating mono-allelic expression of these genes in somatic cells (Figure 1E, F). We found that, silenced *Impact* and *Peg3* alleles became activate during reprogramming, indicating that imprints become erased during iPSC reprogramming. These results are in agreement with previous studies in iPSCs (40, 44) (Figure 1E, F). Thus, in this system, polymorphic female somatic cells with an Xi can be robustly induced to reprogram while enabling allele-resolution gene expression analyses.

### XCR initiates early during entry into pluripotency

We next set out to precisely investigate when reactivation of the Xi takes place. First, we evaluated the expression of the maternally derived X-GFP allele. Transgenic GFP transcripts were detected already at day 10, preceding detection of GFP fluorescence by about 2 days, and following reactivation of several pluripotency TFs (Figure 1G, D). Next, to achieve a better measurement of the erasure of Xi silencing, we calculated the mean expression ratio of genes on the X-chromosome relative to autosomes. We observed chromosome-wide XCR, reflected by a progressive increase in the X-to-autosome (X/A) gene expression ratio in SSEA1+ intermediates starting at day 10 onwards (Figure 1H). As a control, gene expression ratios from chromosome 2 or 8 did not change over time (Figure 1H). These results revealed upregulation of gene expression from the X-chromosome during reprogramming to iPSCs. In order to determine if the increase in X/A gene expression ratio resulted from the reactivation of Xi rather than the upregulation of a single active X-chromosome (Xa), we measured the average allelic expression ratios between *Mus* and *Cast* alleles on the X-chromosome throughout reprogramming (see Methods). Reactivation appeared to start after day 8, and was completed in late reprogramming stages, reaching on average equal expression levels between the two X-chromosomes in late reprogramming intermediates (Figure 1I). By contrast, autosomal genes, on average, maintained similar allelic gene expression ratios throughout reprogramming, confirming increased X-chromosome dosage (Figure S1D). These analyses are consistent with an increase in X-chromosome dosage in iPSCs (45) and robustly showed chromosome-wide erasure of dosage compensation and reactivation of the Xi (Figure 1I).

### Transcriptional reactivation of X-linked genes during induction of pluripotency is gradual and takes several days

To date, the precise chronology of chromosome-wide transcriptional activation during reversal of Xi silencing during iPSC reprogramming has not yet been established. To reveal the dynamics of XCR, we generated time-resolved maps of X-linked allelic expression ratios. Specifically, we calculated the ratio of maternal to total reads for all informative and high confidence X-linked genes using stringent criteria (see Methods). We extracted complete allelic information for 156 X-linked genes. This approach revealed that transcriptional activation of the Xi progresses with gene-specific kinetics (Figure 2A). Surprisingly, several genes (11%, 18/156) were reactivated as early as day 8 of reprogramming, hereafter called ‘early’ reactivated genes (Figure 2A, S2A). This contrasted with a previous studies that reported reactivation of X-linked genes much later in reprogramming, after activation of late pluripotency marker DPPA4 and after complete *Xist* silencing (26, 27). Other genes were delayed in transcriptional reactivation and could be segregated into different groups: with reactivation kinetics between day 8 and day 10 (‘intermediate’), day 10 and day 13 (‘late’) and after day 13 (‘very late’) (Figure 2A). This is in striking contrast with the rapid reversal of the imprinted paternal Xi in the epiblast, which takes about 12 hours (15).

**Figure 2.**
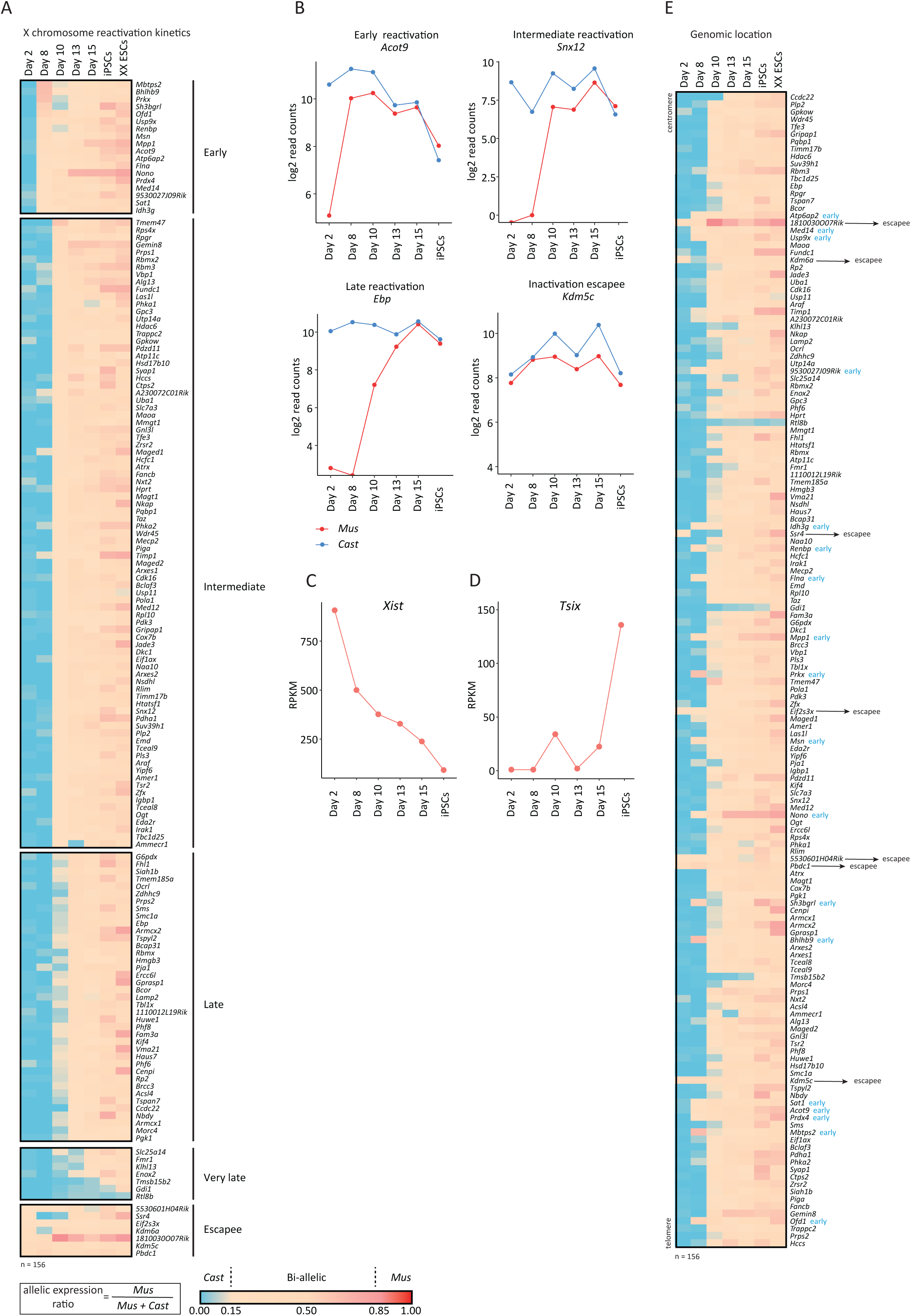
Different X-linked genes reactivate with different kinetics during factor-induced reprogramming to iPSCs. (A) X-linked genes ordered by reactivation timing at different time points of reprogramming within day 2 to 15 and in iPSCs and ESCs. Ratios were calculated by dividing maternal by total reads (*Mus*/*Mus*+*Cast*). The color gradient represents the parental origin of allelic expression, with *Cast* expression in blue (ratio<0.15), *Mus* expression in red (ratio>0.85) and bi-allelic expression range of 0.15-0.85. Number of informative genes =156. Genes were considered early if expressed bi-allelically at day 8, intermediate at day 10, late at day 13, very late at day 15 and escapees at day 2. (B) Gene expression levels (log2 transformed read counts) of representative X-linked genes for each reactivation class (early, intermediate, late and escapee) during reprogramming. Parental origin of the allelic expression is indicated in blue for paternal origin (*Cast*) and in red for maternal origin (*Mus*). (C) *Xist* expression levels (exon to intron ratio RPKM, see methods) during reprogramming. RPKM = Reads Per Kilobase Per Million. (D) *Tsix* expression levels (exon to intron ratio RPKM, see methods) during reprogramming. (E) Gene reactivation kinetics ordered by genomic location of genes on the X chromosome. Ratios were calculated as described in (A).

By plotting the expression of selected paternal and maternal alleles, we observed that X-linked genes such as *Acot9* very rapidly reactivate, illustrating early XCR events (Figure 2B). On the other hand, *Snx12* and *Ebp* displayed delayed activation of the previously inactive allele, requiring additional 2 and 5 days, respectively, to reach bi-allelic expression (Figure 2B). We also recovered the bi-allelic expression of genes that escape XCI in the starting cells, including several known escapee genes (Figure 2B, S2A). In addition, we observed that the maternal X-GFP allele reactivated at day 10 of reprogramming, representing the genes with delayed reactivation kinetics relative to early genes (Figure 1G). Our results reveal that different genes reactivate with different kinetics during XCR induced by iPSC reprogramming, and that XCR is initiated earlier than previously thought, and completed only after several days.

### A subset of genes reactivates before complete *Xist* silencing

Since loss of the lncRNA *Xist* has been reported to be required for XCR (26), we determined its expression kinetics. We observed gradual downregulation of *Xist* starting from day 8, followed by the activation of its antagonist transcript *Tsix* in iPSCs (Figure 2C, D). The downregulation of *Xist* (Figure 2C) suggests that the molecular machinery required to reverse the silenced state of the Xi is triggered as early as at day 8 during reprogramming, but not yet completed. This is in agreement with complete *Xist* silencing taking place after NANOG reactivation (26). Nevertheless, our time course analysis clearly indicated bi-allelic X-linked gene expression by day 8 of reprogramming (Figure 2A, B, S2A). XCR before *Xist* loss has been reported in ICM (15), but not yet during iPSC reprogramming. Our results show that XCR during reprogramming is initiated before complete loss of *Xist* transcript. In addition, reactivation of early genes preceded the activation of the pluripotency-associated gene *Prdm14,* indicating that XCR initiation occurs before the activation of the entire pluripotency network (Figure 2A, 1D). Altogether, these results indicate that reactivation of different X-linked genes might be mediated by different mechanisms and, for some genes, before the complete loss of *Xist* or full activation of the pluripotency network.

### Gene reactivation kinetics relate to genomic and epigenomic features

We sought to identify features that help to explain the precise timing of X-linked gene reactivation. We first investigated whether there is a link between reactivation kinetics and location on the X-chromosome. To this end, we generated heatmaps of allelic expression and ordered the genes according to their genomic location (Figure 2E). Early genes were distributed throughout the chromosome. Interestingly, we detected two clusters of early reactivated genes. The first cluster contained *Atp6ap7*, *Med14* and *Usp9x*. The second cluster contained *Sat1*, Acot9 and Prdx4.

The timing of transcriptional activation could not be explained by genomic distance to the *Xist* locus (Figure S3A). Moreover, there was no correlation between the timing of gene activation and the level of gene expression in ESCs (Figure S3A). Next, we compared the timing of gene activation during reversal of rXCI with reactivation of iXCI (15). Most of the genes that reactivate early or late were different in both systems (Figure S3B). Hence, the kinetics of XCR during reversal of rXCI are different from that of iXCI reprogramming. Taken together, these results suggest that the reversal of rXCI differs from that of iXCI, and is independent of the genomic distance of genes to the *Xist* locus.

Next, we determined whether the timing of reactivation could be associated with the presence or absence of chromatin marks on the starting Xi. We used allele-specific chromatin immunoprecipitation sequencing (ChIP*-*seq) data for H3K27me3, H3K36me3 and H3K4me3 in MEFs (46) to determine the enrichment of chromatin marks around the transcription starting sites (TSS) of early, intermediate, late and very late reactivated genes. We found that on the Xi, H3K27me3 is clearly enriched on all classes of genes, but there were no clear differences between classes, except for escapee genes which displayed much lower levels of H3K27me3 (escapee vs intermediate *p*=0.016, escapee vs late *p*=0.045 by Wilcoxon rank test). Genes on the Xa had significantly lower levels of H3K27me3 enrichment (Figure S3C). Thus, while H3K27me3 may restrict the activation of silenced genes on the Xi, it is not sufficient to explain the delay in reactivation of late genes. Additionally, on the Xi, H3K36me3 and H4K3me3 were depleted on all classes of genes, but enriched on escapee genes (H3K36me3: escapee vs early *p*=0.02; escapee vs intermediate *p*=0.005; escapee vs late *p*=0.003; H3K4me3: escapees vs early *p*=0.039; escapees vs intermediate *p*=0.0035; escapees vs late *p*=0.002 by Wilcoxon rank test) (Figure S3C). On the Xa, early reactivating genes were more enriched in H3K36me3, but not H3K4me3, compared to the late ones. Altogether, this analysis revealed that different classes of genes do not appear to show significant differences in the chromatin marks examined on the Xi in MEFs. Therefore, the timing of XCR cannot be fully explained by different levels of these histone modifications on the Xi.

During XCI, silenced genes relocalize to the interior of the repressive Xi compartment, while escapee genes remain at the periphery of the *Xist* domain, outside of the silenced compartment occupied by the Xi. This allows them to avoid the silencing machinery (47-53). Hence, we asked whether early reactivated genes are located closer to escapee genes, where they might reside in topologically favorable positions that facilitate their reactivation during reprogramming. To address this, we measured the average genomic distance between the different classes of genes and the nearest escapee gene. We found a clear and significant decrease in the genomic distance to nearest escapees for early and for intermediate genes compared to very late reactivated genes (respectively *p*=0.021 and *p*=0.038 by Wilcoxon rank test) (Figure 3A). Moreover, the two clusters of early genes each had a nearby escapee gene in the genomic sequence map (Figure 2E). Altogether, these results reveal that genes that reactivate early during iPSC reprogramming have significantly reduced genomic distance to escapee genes compared to very late genes. Overall, this suggests that the localization of a gene in 3D space relative to repressive chromatin domains may be related to the stability of gene silencing.

**Figure 3.**
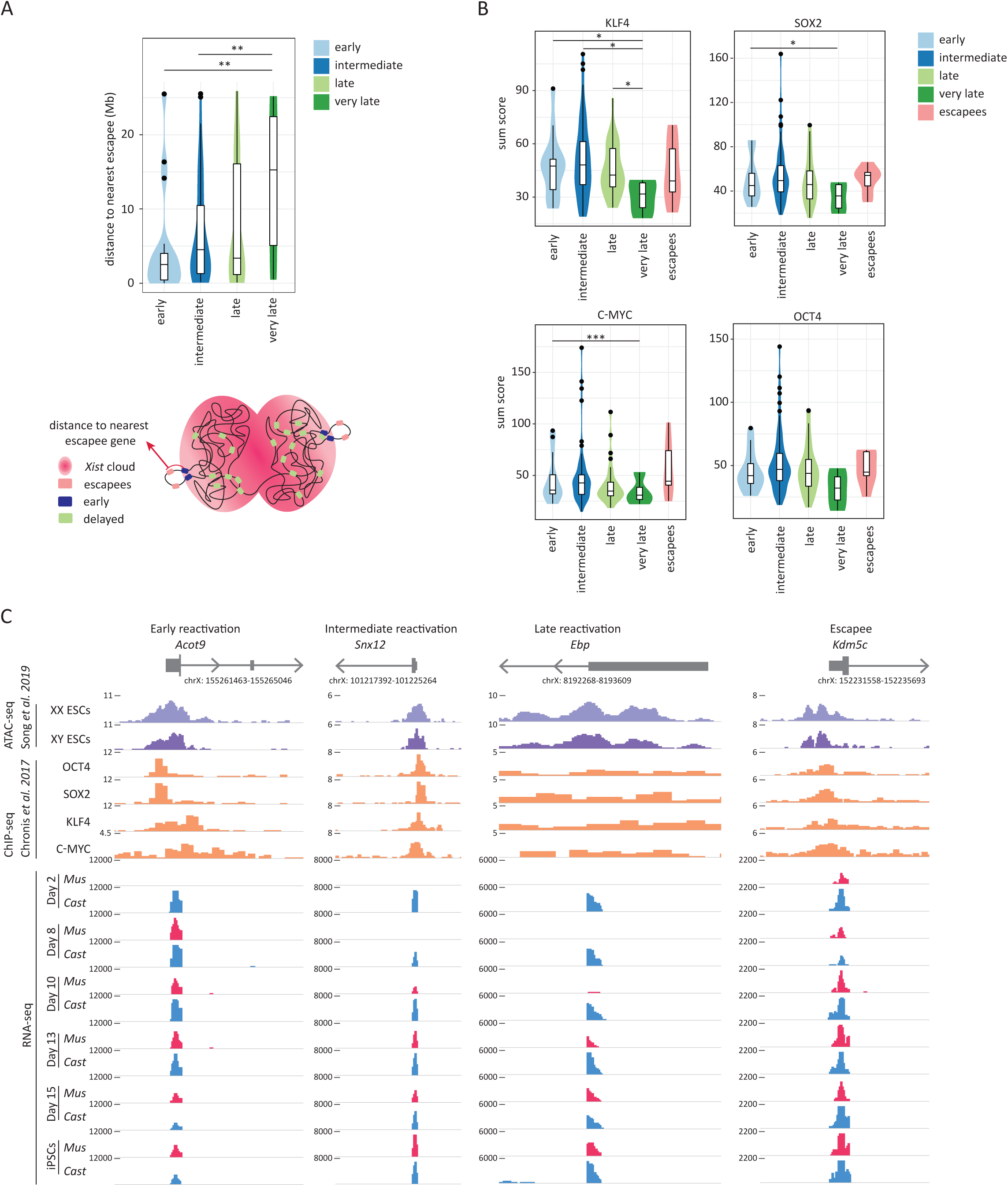
Link between genomic and epigenomic features and XCR kinetics. (A) Violin diagram with the distance to nearest escapee (Mb) for each reactivation class. Below by schematic figure of distance to nearest escapee gene. The significant *p*- values of a Wilcoxon rank test comparing the different gene reactivation classes are indicated with asterisks above the violin plots (****, *p*- value=0.0001-0.001=extremely significant; ***, 0.001-0.01=very significant; **, *p*-value=0.01-0.05=significant; *, *p*- value≥0.05=not significant). (B) Violin plots indicating the sum score of enrichment levels of OCT4, SOX2, KLF4 and C-MYC occupancy for each reactivation class (early, intermediate, late, very late and escapees). The *p*- values are calculated as described above. (C) ATAC-seq signal for open chromatin in female and male ESCs, ChIP-seq signal for OCT4, SOX2, KLF4 and C-MYC binding in male ESCs and RNA-seq signal of allele resolution gene expression during reprogramming (blue for *Cast* origin and red for *Mus* origin) for a representative gene corresponding to each reactivation class (early, intermediate, late and escapee).

### The timing of gene reactivation is linked with pluripotency TF binding

TFs are primary mediators of gene regulatory programs during development and their binding to cis-regulatory elements is often followed by or concomitant with reorganization of chromatin, the displacement of repressive modifications and regulators, and the recruitment of active ones leading to transcriptional activity (54). It remains unclear how TFs mediate XCR during reprogramming and in particular if they may contribute to the differential reactivation kinetics of different X-linked genes. To address these questions, we set out to investigate whether differently reactivating genes have distinct enrichment of pluripotency TFs binding. To examine the enrichment levels of pluripotency TFs at the TSS of reactivating genes, we used previously published ChIP*-*seq dataset with binding profiles of four reprogramming TFs (OCT4, SOX2, KLF4, C-MYC) on the Xa in male ESCs (55) (Figure 3B). We found that early genes showed a significantly higher enrichment of KLF4, SOX2 and C-MYC binding compared to very late reactivating genes (Figure 3B, *p*=0,046; *p*=0,025; and *p=*0,0078 respectively by Wilcoxon test). Intermediate and late reactivating genes also showed a significant increase in the enrichment of KLF4 compared to very late genes. Thus, Xi-linked genes might be targeted by pluripotency TFs for their transcriptional activation in iPSCs.

We next visualized TF enrichment at the TSS of *Acot9* (early), *Snx12* (intermediate), *Ebp* (late) and *Kdm5c* (escapee). In iPSCs, these genes are bi-allelically expressed and have accessible chromatin at their TSS in ESCs (Figure 3C). We found a clear enrichment of several pluripotency TFs binding at the TSS of *Acot9*, *Snx12* and *Kdm5c*, but not late reactivated gene *Ebp*. These observations strengthen the possibility that pluripotency TFs might directly activate X-linked gene expression in iPSCs. Next, we asked whether pluripotency TFs possess the capacity to bind X-linked genes during the reprogramming process. We analyzed ChIP*-*seq data for SOX2 and OCT4 in male SSEA1+ reprogramming intermediates, because such data is unfortunately not yet available in female cells (56). Nevertheless, we found that several X-linked genes showed enrichment of SOX2 and OCT4 binding at their promoters in reprogramming intermediates (Figure S3D). We conclude that pluripotency TFs such as SOX2 and OCT4 possess the ability to bind X-linked genes, at least on the Xa, during the reprogramming process. Therefore, pluripotency TFs may target early and intermediate genes more efficiently than late genes during the acquisition of the iPSC state.

In conclusion, the early reactivation of genes during entry into pluripotency and before complete *Xist* loss may be related to the distance of these genes to the nearest escapee gene, and their associated topology, as well as preferential targeting by pluripotency TFs. Most of the very late genes, however, may be embedded in the more repressed compartment of the Xi that is coated by *Xist* lncRNA, and less efficiently targeted by pluripotency TFs. The precise role of chromatin topology, chromatin accessibility and TF binding in relation to chromosome-wide reversal of gene silencing merits further exploration.

### XCR is restricted by the removal of active histone marks during reprogramming to iPSCs

TF binding to gene regulatory elements mediates changes in chromatin. Hence, we aimed to identify chromatin pathways that functionally restrict or facilitate reversal of XCI during entry into the iPSC state. We used the X-GFP reporter to test whether interference with different chromatin pathways has an effect on transcriptional activation of non-early genes (Figure 1G). We induced reprogramming of female MEFs with Xi-GFP into iPSCs and carried out an epigenetic drug screen in order to identify chromatin regulators that might act as barriers or mediators of XCR (Figure 4A). We included drugs that target several factors involved in chromatin regulation, such as DNA methylation, repressive histone modifications as well as in the regulation of DNA topology and chromatin remodeling (Figure 4B) (57-59). After 10 days of reprogramming and continuous treatment with epigenetic drugs, we recorded the proportion of cells that activated SSEA1 and GFP expression. This enabled us to define the effects of drug treatment on XCR in cells undergoing productive reprogramming (SSEA1+ cells) versus non-productive reprogramming intermediates (SSEA1-).

**Figure 4.**
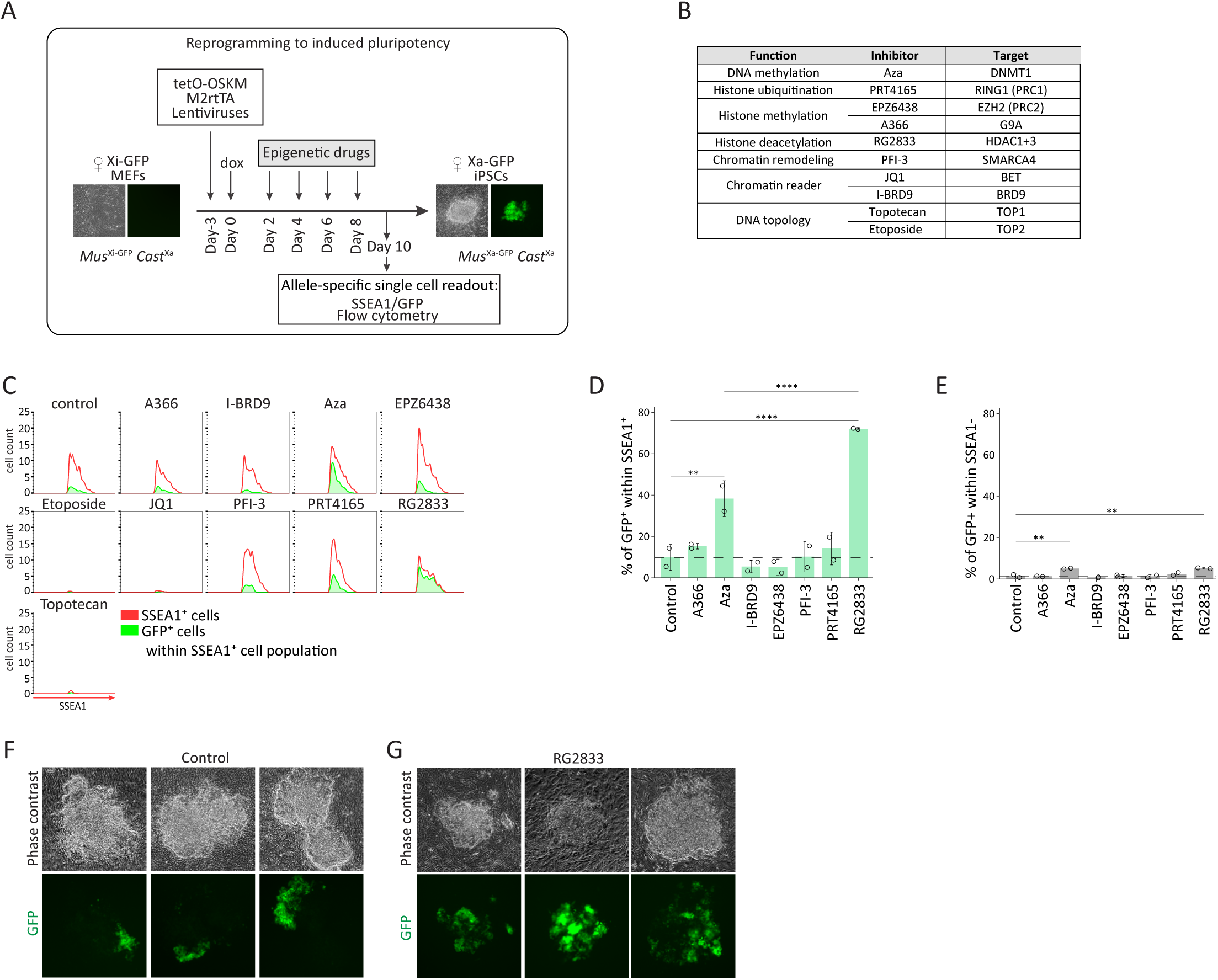
Histone deacetylases restrict XCR during reprogramming to iPSCs. (A) A scheme representing experimental design of epigenetic drug inhibitor screen during reprogramming. (B) Inhibitors added individually at different time points during reprogramming with their function, name and target molecule(s). (C) Histograms representing the flow cytometry analysis of the proportion of GFP+ cells within SSEA1+ cell for each individual inhibitor at day 10 of reprogramming and vehicle control. The whole SSEA1+ cell population is represented in red. In green, population of GFP+ cells within SSEA1+ cells. (D) Proportion of GFP+ cells within the SSEA1+ cell population for each individual inhibitor. The *p*-values of one-way ANOVA with Dunnett’s multiple comparisons test comparing levels of GFP+ cells after treatment with each individual inhibitor and vehicle control are indicated with asterisks above the violin plots. The *p*-values of one-way ANOVA with Sidak’s multiple comparisons test comparing levels of GFP+ cells treated with RG2833 and Aza are indicated with asterisks above the boxplot (****, *p*-value=0.0001-0.001=extremely significant; ***, 0.001-0.01=very significant; **, *p*-value=0.01-0.05=significant; *, *p*-value≥0.05=not significant). n = 2. (E) Proportion of GFP+ cells within SSEA1-cell population for each individual inhibitor. The *p*-values of one-way ANOVA with Dunnett’s multiple comparisons test comparing levels of GFP-cells after treatment with each individual drug inhibitor and vehicle control are indicated with asterisks above the boxplot. n = 2. (F) Phase contrast and fluorescent images of three representative colonies at day 10 of reprogramming after treatment with DMSO control during epigenetic drug screening. (G) Phase contrast and fluorescent images of three representative colonies at day 10 of reprogramming after treatment with RG2833, an inhibitor of HDAC1/3, during epigenetic drug screening.

We found that treatment with RG2833 or Aza both led to a significant increase in XCR within SSEA1+ cells relative to untreated control (Figure 4C, D, S4B). RG2833 is an HDAC3 and HDAC1 inhibitor, while Aza inhibits the maintenance DNA methyltransferase DNMT1. The proportion of GFP+ cells after treatment with RG2833 and Aza was much higher in SSEA1+ than in SSEA1-cells, distinguishing effects of XCR in reprogramming versus loss of XCI maintenance in somatic cells (Figure 4E). The effects of RG2833 were also clearly visible when following the reactivation of X-GFP transgene in iPSCs colonies after 10 days of reprogramming (Figure 4F, G). The increase in XCR upon Aza and RG2833 treatment was not the consequence of an increase in reprogramming efficiency, because we did not detect significant differences in the number of SSEA1+ cells between conditions (Figure S4A). These results corroborated a previous study in which DNA methylation was reported to oppose XCR during the generation of iPSCs (26). Thus, our screen recovered a known barrier to XCR, but also identified HDACs as potential barriers to XCR during pluripotency induction, which has not previously been implicated. Interestingly, recent work from Zylicz *et al*. revealed that HDAC3 activation on the X-chromosome initiates transcriptional silencing during initiation of XCI (1). Therefore, recruitment of HDAC3, through *Xist* to the Xi, and of HDAC1, may deacetylate silenced genes to oppose transcriptional activation on the Xi, acting as a barrier to XCR during iPSC reprogramming. We conclude that several chromatin pathways, including histone deacetylation, oppose XCR in reprogramming intermediates.

### Chromatin acetylation on Xi is prevented until late reprogramming stages

The findings above indicate that deacetylation of histones might prevent timely reactivation of X-linked genes. To explore the dynamics of histone acetylation of the Xi during reprogramming to induced pluripotency, we performed time course immunofluorescence analysis throughout reprogramming for H3K27ac, H3K27me3 and NANOG. In the starting MEFs, we found that the Xi, marked by H3K27me3 enrichment, was depleted of H3K27ac mark (Figure 5A). During reprogramming, the hypoacetylated state of the Xi was maintained in NANOG+ cells when H3K27me3 was still present and NANOG was excluded from the Xi (Figure 5A-C, S5A). Established iPSCs lost both H3K27me3 enrichment and gained H3K27ac and NANOG, in agreement with late global acetylation of the Xi during XCR. These results suggest that histone hypoacetylation on the Xi persists until late stages of XCR and may be maintained by the constant action of histone deacetylases. We conclude that changes in chromatin states during XCR involve chromatin acetylation, at a time when TFs may engage with chromatin of the Xi leading to transcriptional activation. Altogether, our results provide a broader understanding of how TFs induce dynamic reversal of stable transcriptional silencing and overcome active barriers to transcriptional activation.

**Figure 5.**
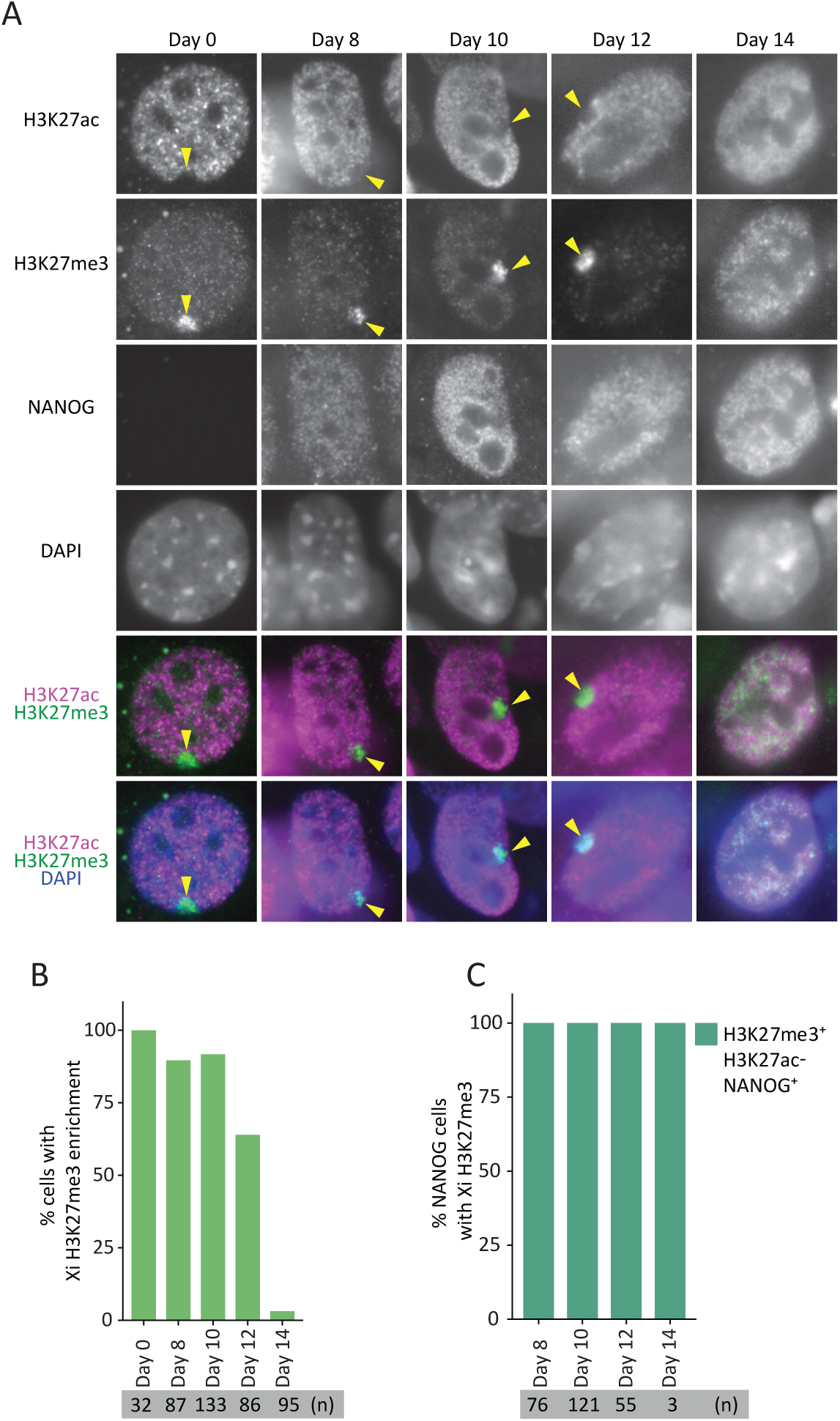
Hypoacetylation of Xi persists until late stages of reprogramming. (A) Immunofluorescence analysis for H3K27ac (magenta in merge), H3K27me3 (green) and NANOG at different stages of reprogramming. Dapi staining (blue) marks nuclei. Xi H3K27me3 enrichment is marked with an arrowhead. (B) Proportion of cells with H3K27me3 enrichment during reprogramming. (C) Proportion of NANOG+ cells with H3K27me3 enrichment and H3K27 hypoacetylation during reprogramming.

## DISCUSSION

Reversal of epigenetic memory on the Xi during pluripotency induction is a paradigm for studying transcriptional activation of silenced chromatin. However, chromosome-wide allelic gene activation has not yet been completely grasped. Furthermore, the relationship between pluripotency TFs and reversal of gene silencing has remained largely underexplored. We used transcriptomic and epigenomic approaches to define allele-resolution maps of chromosome-wide gene activation during reprogramming to iPSCs. This allowed us to focus on the exploration of the progressive nature of XCI reversal in reprogramming, where different genes seem to exhibit distinct levels of silencing stability and as a result reactivate with different timing (Figure 6). We found that gene activation is initiated before the upregulation of late pluripotency genes such as *Prdm14* and prior to complete silencing of the lncRNA *Xist*, but is completed late during reprogramming (Figure 6). We then interrogated the relationship between the timing of transcriptional activation and genomic and epigenomic features. We showed that neither the distance of X-linked genes to the *Xist* locus, gene expression nor enrichment of H3K27me3 on Xi can explain different reactivation kinetics. We revealed that early reactivating genes tend to reside in regions genomically closer to genes that escape XCI and might be preferentially targeted by pluripotency TFs (Figure 6). Furthermore, to better understand the mechanisms underlying the prolonged nature of XCR during reprogramming, we employed epigenetic drug screens and identified histone deacetylation as a barrier to gene reactivation from the Xi (Figure 6). Altogether, we provide a framework for how TFs induce dynamic chromosome-wide reversal of stable epigenetic memory by overcoming active barriers in order to activate gene expression.

**Figure 6.**
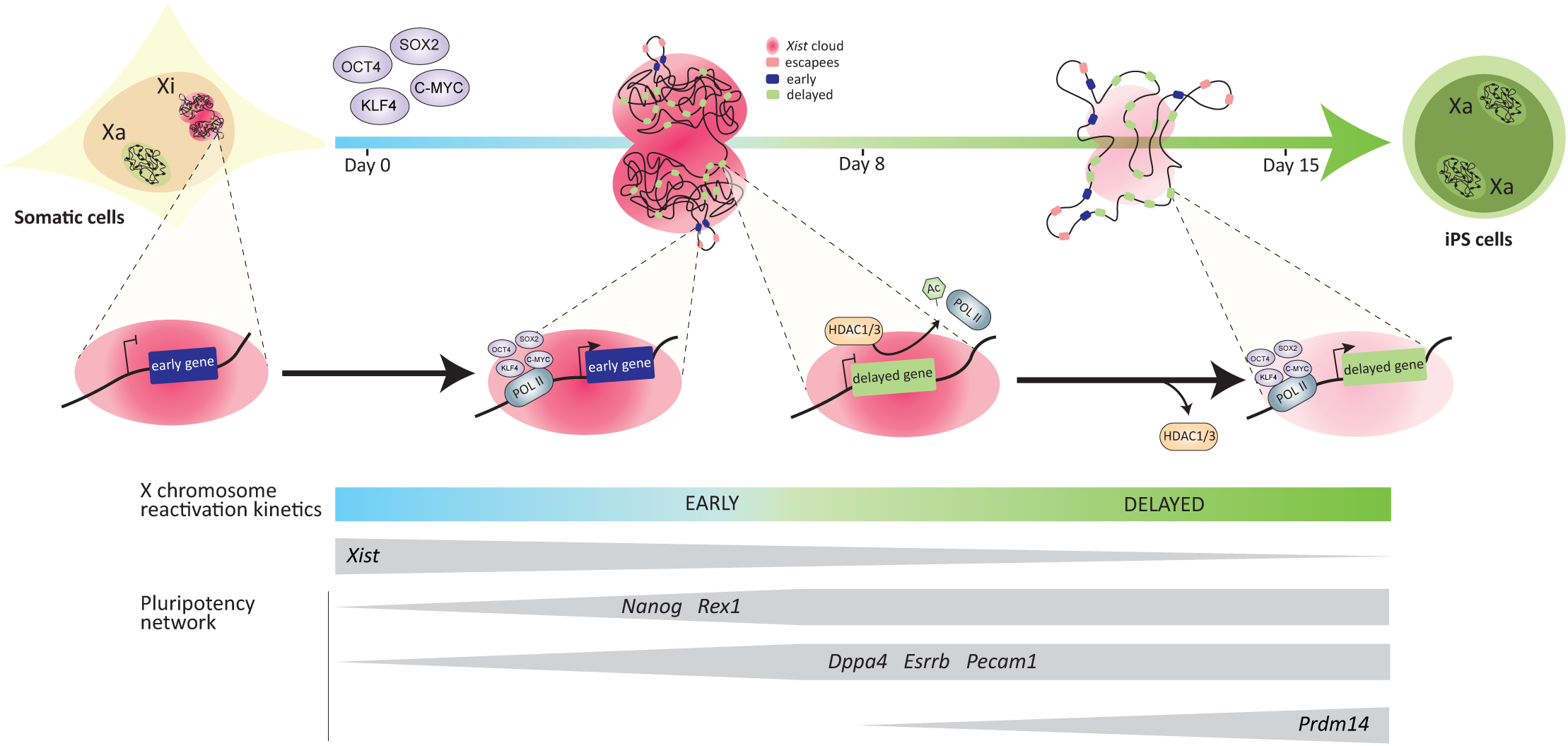
Dynamics of XCR during somatic cell reprogramming to iPSCs. In starting female somatic cells, Xi is coated by the lncRNA *Xist*. Reactivation of early genes from the Xi initiates at early onset of reprogramming, preceding full repression of *Xist* and the establishment of complete pluripotency network. Those early genes are located linearly closer to escapee genes, hence might reside in topologically favorable position. In addition, early reactivating genes might be preferentially targeted by pluripotency TFs. Upon further upregulation of pluripotency network and downregulation of *Xist*, other X-linked genes reactivate with hierarchical kinetics for several more days. The prolonged nature of XCR might be attributed to the action of HDAC1/3 which restrict reactivation of X-linked genes during reprogramming, possibly by maintaining high hypoacetylation state of the Xi.

### XCR is rapidly initiated during entry into pluripotency

We found that the machinery to induce silencing reversal starts to act early during the onset of reprogramming to iPSCs. We show that the lncRNA *Xist* is downregulated already at the entry into pluripotency when TFs such as *Nanog* become first expressed. The early initiation of XCR is in contrast with a previous report that XCR takes place very late during reprogramming to iPSCs (26). The differences might be due to the use of allele-specific RNA-seq compared to the previously used RNA-FISH. Many studies indicated a role of pluripotency TFs in XCR induction (25, 33, 36). A robust naïve pluripotency network in mouse ESCs has been shown to be required for complete suppression of *Xist* (33). However, the mechanism through which these factors silence *Xist* is still unclear (36). Our data suggest that pluripotency TFs may also be able to directly bind to regulatory elements of many X-linked genes during reprogramming and in ESCs. Although there might be different mechanisms at play, it is possible that pluripotency TFs act as pioneer factors opening up chromatin on the Xi or suppress the silencing effect of the nuclear lamina where Xi is localized (60). Allele-resolution chromatin accessibility analyses could help further address these questions. The precise molecular mechanisms by which *Xist* downregulation and XCR induction take place remain intriguing. Additionally, the initiation of XCR during reprogramming is challenging to capture due to the degree of heterogeneity associated with factor-induced iPSC reprogramming. To minimize the variability, our study provides information from SSEA1 sorted cells that have been shown as a robust marker of cells poised to reprogram successfully (38). Nevertheless, single cell allele-resolution approaches would allow to pinpoint most upstream events of XCR.

### A subset of X-linked genes reactivates before complete silencing of *Xist* and before pluripotency is fully established

In line with early XCR initiation, we found that a subset of genes reactivates early and before complete loss of *Xist* and the acquisition of a complete pluripotency network. This is in agreement with studies of iXCI reactivation (15). However, we found that reactivation of rXCI differs from that of iXCI in terms of the timing necessary to complete reactivation as well as the order of reactivating genes. This may be explained by different starting epigenetic states. Nevertheless, early reactivation in both cases suggests a specific mechanism that allows genes to bypass *Xist’s* activity and reactivate rapidly. We hypothesize that the 3D architecture of the Xi might play an important role (8). It is possible that genes closer to borders of the silent compartment might be more easily unfolded and exposed to TFs. Indeed, we show that the linear genomic distance of early reactivating genes to escapees, known to reside outside of megadomains (9), is shorter. This opens the exciting possibility that the order of conformational changes might be causally linked with reactivation timing, meriting further exploration.

### Many X-linked genes reactivate with a significant delay

Our data also shows the protracted nature of XCR for most X-linked genes. After reactivation of early genes, a significant amount of time is required to complete activation of other loci. This suggests a possible mechanism where suppressed chromatin has to be first remodeled in order to allow subsequent gene activation. We show however that the accumulation of histone marks such as H3K27me3 on the Xi of MEFs is insufficient to explain the delay in reactivation. Neither enrichment of active marks such as H3K4me3 and H3K36me3 can support early reactivation. This is in contrast to reactivation of iXCI where H3K27me3 enrichment on Xi had a substantial impact on reactivation dynamics (15). It is possible that during reactivation of rXCI, other repressive marks such as macroH2A, H2AK119Ub or DNA methylation are at play. Our study indicates that pluripotency TFs that accumulate during reprogramming to induced pluripotency might play an important role in dictating gene activation kinetics. We found that on the Xa in ESCs, four pluripotency factors (OCT4, SOX2, KLF4 and C-MYC) differentially bind to early and non-early reactivating genes. KLF4, C-MYC and SOX2 show significantly lower level of enrichment around cis regulatory regions of very late reactivating genes compared to early ones. This suggests that genes reactivating with the highest degree of delay might be less accessible to those TFs. The possible reasons for that, as discussed above, might be linked to three dimensional topology of the X-chromosome, chromatin accessibility or other repressive features that obstruct TFs’ binding sites. Allele-resolution HiC, chromatin accessibility and ChIP-seq analyses will be needed for further clarification.

### Histone deacetylases oppose rapid XCR

Here, using epigenetic drug screening with a GFP reporter for activation of delayed genes, we identified histone deacetylases as barriers to reactivation that contribute to the delayed nature of XCR. It has been recently shown that histone deacetylation is the most efficiently engaged suppressive chromatin mechanism during initiation of XCI (1), but the role of histone deacetylation in XCR during iPSC reprogramming was not known. We show that the inhibition of HDACs accelerates reactivation and thus highlights an important mechanism of silencing stability. The link between transcriptional activity and histone acetylation has been very well established and shown to be conserved throughout a wide range of species (61). In the context of XCI, synergistic effects of *Xist* coating, methylation of CpG islands and hypoacetylation of histone H3 and H4 have been reported as features associated with the establishment and maintenance of the Xi in somatic cells (62). Without the induction of pluripotency, however, inhibition of HDACs does not lead to complete reactivation likely because DNA methylation and *Xist* still stabilize Xi (57). It is possible that after XCI, HDACs remain bound to safeguard Xi silencing. Histone acetylation and deacetylation establish a regulatory balance, which changes depending on gene activity (61). We show here that the hypoacetylated state of the Xi persists until very late stages of the reprogramming. Whether HDACs are recruited continuously or whether histone acetyltransferases are unable to acetylate residues remains unclear.

## CONCLUSION

Despite the stable epigenetic memory of random XCI, transcriptional silencing can be gradually reversed by reprogramming to iPSCs. The genes that are transcriptionally activated early tend to reside genomically closer to escapee genes, and may be preferentially targeted by pluripotency TFs for their reactivation. Additionally, XCR during the iniduction to pluripotency is restricted by several chromatin pathways including histone deacetylation. In sum, our findings reveal relationships between genomic and epigenomic features on the one hand and the reversal of gene silencing during cell fate reprogramming on the other hand. Our results open up avenues for a better understanding of allelic gene regulation and epigenetic reprogramming.

## EXPERIMENTAL PROCEDURES

### Cell lines

X-GFP reporter MEFs were derived from female E13.5 embryos hemizygous for the X-GFP transgenic allele (63). These embryos resulted from the cross between female X-GFP *Mus musculus musculus* and male *Mus musculus castaneus* lines (37). Stem cell cassette (STEMCCA) MEFs refers to female MEFs carrying a single polycistronic reprogramming cassette with four reprogramming factors OSKM (referred to as tetO-OSKM) located in the *Col1A* locus together with a single copy of reverse tetracycline transcactivator M2rtTA in the Rosa26 locus. iPSC controls were derived from reprogramming experiments in this study described below. ESC-like colonies were picked at day 14 of reprogramming and cultured for four passages in conditions described below.

### Cell culture and reprogramming methods

MEFs were cultured in MEF medium [DMEM (Gibco, 41966-554 052) supplemented with 10% (v/v) fetal bovine serum (FBS, Gibco, 10270-106), 1% (v/v) penicillin/streptomycin (P/S, Gibco, 15140-122), 1% (v/v) GlutaMAX (Gibco, 35050-061), 1% (v/v) non-essential amino acids (NEAA, Gibco, 11140-050), and 0.008% (v/v) beta-mercaptoethanol (Sigma, M7522)].

Reprogramming experiments, unless stated otherwise, were performed by conditional induction of lentivirally delivered reprogramming factors. First, P1 MEFs at around 70% confluency were transduced with concentrated lentiviral supernatants. Lentiviruses were generated using HEK cells separately for two constructs: tetO-FUW-OSKM, Addgene cat. 20321 (64) and FUW-M2rtTA, Addgene cat. 20342 (65) with the calcium precipitation method. Supernatants with lentiviral particles were concentrated using lenti-X-concentrator (Takara, 631231) in 1:100 ratios. Infection with a pool of equal volumes of both constructs was carried out overnight, followed by 12 hours culture in MEF medium and 1:5 split. Cells then were sorted using FACS (described below) in order to isolate homogeneous population with regard to allelic inactivation of the X-GFP transgene (either Xi-GFP or Xa-GFP). Cells were plated for reprogramming directly after sorting, 50,000 cells per one well of a 12 well plate. Reprogramming was induced by doxycycline (2 μg/ml final) in mouse ESC medium [KnockOut DMEM (Gibco, 10829-018) supplemented with 15% FBS, 1% P/S 10,000 U/mL, 1% GlutaMAX 100X, 1% NEAA 100X, 0.008% (v/v) beta-mercaptoethanol, and mouse LIF] in the presence of ascorbic acid (50 μg/ml final). The medium together with doxycycline and ascorbic acid was replaced every two days and maintained throughout entire reprogramming experiments. To derive iPSCs from picked colonies, the same conditions were used with dox and AA withdrawal after picking. STEMCCA reprogramming was carried out as previously reported (26).

### Immunofluorescence

Immunofluorescence staining was carried out as previously described (26). The primary antibodies used were as follows: H3K27me3 (mouse, 1:500, Active motif, cat. 61017), H3K27ac (rabbit, 1:500, Abcam, cat. Ab4729), NANOG (rat, 1:200, eBioscence, cat. 14-5761-80). Images were captured with a Zeiss Axioimager Z1 inverted microscope coupled with an AxioCam MRc5 camera and Axio Vision software. Multi-channel images were cropped and merged in ImageJ. The number of scored cells are indicated under each plot.

### Flow cytometry and cell sorting

For cell sorting and flow cytometry analysis cells were dissociated using trypsin digestion. For X-GFP+/-cell sorting dissociation was followed by washing in the incubation buffer (1x PBS, 0.5% BSA, 2mM EDTA) and filtered through Falcon^®^ 40 µm Cell Strainer (Corning, cat. 352340). For sorting SSEA1+ cells as well as for flow cytometry analysis in screening approaches, dissociation was followed by wash in the incubation buffer and 40 minutes of incubation with primary antibody anti SSEA1 coupled with PE (Mouse IgM, R&D, FAB2155P, Clone MC-480, conc. 1 µl SSEA1-PE Ab / 5.10^6^ cells). Stained cells were subsequently washed in incubation buffer to remove residual unbound antibody and passed through a cell strainer. Cell death exclusion was always applied by staining with DAPI (Sigma cat. D9542-50MG). Sorting was performed on a BD FACS Aria III or BD Influx (BD Biosciences) and performed by expert operators at KU Leuven FACS core. Flow cytometry was performed on BD Canto II HTS.

### RNA-seq library preparation

RNA-seq library was prepared from low RNA input using an adapted Smart-seq2 protocol (66). Briefly, sorted SSEA1+ reprogramming intermediates were immediately lysed in RLT buffer (RNeasy Micro Kit (Qiagen, 74004)) and stored in −80°C until cells from all timepoints were collected. Next, all samples were processed together to extract RNA following the manufacturer’s protocol. cDNA synthesis was done starting from 500 pg of input RNA, followed by library preparation from 80 pg of cDNA using Nextera XT kit (Illumina, FC-131-1096). Indexing was performed with the Nextera XT index Kit V2 (Illumina, FC-131-2003). The quality of input RNA, cDNA and individual libraries was assessed using a Bioanalyzer (Agilent 2100 Bioanalyzer system). Libraries were pooled and sequenced at the VIB Nucleomics Core on a NextSeq500 (Illumina) sequencer in high-output paired-end mode (2×75 bp) yielding on average 66 million reads per sample (details: Supplementary Table 1).

### RNA-seq reads processing

Read quality was initially assessed using FastQC (94% >Q30). For non-allele resolution analyses, reads were aligned to the mouse reference genome (mm10; GRCm38.p5) using STAR 2.5.3a supplied with the corresponding gencode vM16 annotation file. Mapping was done with default parameters with specified --sjdbOverhang 74 followed by conversion to sorted BAM files. On average, 83.96% of reads were uniquely mapped and only those were passed to next steps. Subsequently, the featureCounts function from the R Bioconductor package “Rsubread” (version 1.5.2) was used to assign mapped reads to genomic features. For allele resolution analyses, reads were mapped to the same reference genome release in which SNP positions were substituted by N base (referred to as N-masked mm10). N-masking was performed with SNPsplit software (Version 0.3.2 released (29-03-2017)) supplied with the list of strain specific SNPs (129S1_SvImJ and CAST_EiJ) from Sanger Mouse Genomes project database (mgp.v5.merged.snps_all.dbSNP142.vcf.gz). N-masking was done in dual hybrid mode and resulted in the identification throughout entire genome of 20,563,466 SNP positions unique for either strain, of which 634,730 on the X-chromosome. Next, reads were aligned to the N-masked mm10 genome using STAR 2.5.3a with parameters disabling soft-clipping of incompletely aligned reads (--alignEndsType EndToEnd --outSAMattributes NH HI NM MD). Reads aligned to the N-masked reference genome were then splitted into two BAM files containing only strain-specific reads (on average 10.95% for *Mus* and 10.25% for *Cast* of total mapped reads) using SNPsplit (details: Supplementary Table 2). Furthermore, to create count matrices with only reliable allelic information, only SNPs covered by at least 5 reads were selected and genes comprising of at least 8 of such SNPs were retained (details: Supplementary Table 3).

### PCA and gene expression analysis

Processing raw read counts for non-allele-specific analysis was supported by the DESeq2 package and associated protocol (67). Genes that did not express at least 10 reads in total across all libraries were discarded from further analyses. Next, PCA was plotted using plotPCA function from the DESeq2 package with input of top 500 most variable genes after rlog transformation. Unless mentioned otherwise, gene expression was presented as log2 values after size-factor normalization for the differences in library size (DESeq2). To evaluate *Xist* and *Tsix* expression, read coverage was first calculated and normalized for library size (RPKM) using bamCoverage function from deeptools (2.4.1) package with --binsize 1. Next, to overcome lack of strand-specific information, ratio of exon to intron coverage at the exon:intron downstream boundary (100 bp overlap) was calculated at exon 5 and exon 1 for *Xist* and *Tsix,* respectively. For X-chromosome to autosome ratio, the mean expression of all X-linked genes (and as control chromosome 8 and chromosome 2 genes) was divided by the mean expression of all autosomal genes.

### Reactivation dynamics

For the calculation of allelic ratio in Figure 1I, 2A, 2E and S2A, two additional filtering criteria were imposed. First, for each gene at a given time point, the sum of reads from both alleles had to be at least 40. Secondly, in order to unambiguously determine reactivation timing, only the genes that passed previous criterion across all time points were included. In Figure 1I, the ratio was calculated as the log2 ratio of *Mus* allele to *Cast* allele calculated for each gene. In figures 2A, 2E, the allelic ratio is represented as ratio of *Mus* to total: (*Mus*/(*Mus*+*Cast*). The heatmap was generated in the online software Morpheus (Morpheus, https://software.broadinstitute.org/morpheus).

### Allele-specific ChIP-seq data analysis

For TF and histone mark enrichment analyses in regulatory regions of genes with different timing of reactivation, published ChIP-seq data were re-analyzed. The datasets used include: (GSE GSE90893) (55) – pluripotency factors and chromatin marks in ESCs and MEFs, (GSE GSE36905) (46) – histone marks in MEFs with allele-resolution. Raw ChIP-seq data were analyzed using the ChIP-seq pipeline from the Kundaje lab (version 0.3.0). Briefly, raw reads were mapped to the reference genome (mm10) using BWA (version 0.7.13) with dynamic read trimming –q 5 and sorted using samtools (1.2). Unmapped reads, reads with quality <30 and PCR duplicates were discarded. PCR duplicates were marked using Picard (v 1.126). Read coverage was first calculated and normalized for library size (RPKM) using bamCoverage function from deeptools (2.4.1) package with --binsize 100. For allele-specific data processing of Pinter et al 2012 data set, the Kundaje pipeline has been adapted to accommodate the SNPsplit pipeline (described above). Briefly, additional parameter has been set for BWA alignment (-s in sampe function) to prevent soft-clipping. Next, filtered BAM files were processed with SNPsplit and the Kundaje pipeline was continued as described above.

To assess the enrichment of selected marks or TFs, first TSSs of the genes with defined reactivation kinetics were selected. Where there were more than one TSS per gene, the first upstream TSS was selected. Enrichment values were calculated by summing the score within 5 kb around the TSS using bedops (2.4.35). Statistical significance of differences between enrichment levels in different reactivation classes was measured using Wilcoxon rank test.

### Epidrug screen

Epigenetic inhibitors were obtained from Selleckchem or Tocris and added to mouse ESC medium in concentrations as previously described (57) (details: Supplementary Table 4). Cells were treated with inhibitors starting from day 2 during reprogramming and inhibitors were refreshed every other day by media change. To calculate the statistical significance of differences, one-way ANOVA with Dunnett’s multiple comparisons test was used to compare the effect of each individual treatment with vehicle control. One-way ANOVA with Sidak’s multiple comparisons test was used to determine significance level between the effect of treatment with RG2833 and Aza.

### Live imaging

Images representing the activity of the X-GFP reporter and phase contrast were acquired using a Nikon Eclipse Ti2 microscope coupled with Nikon NIS Elements software. Images were exported using NIS Viewer software and cropped in ImageJ.

## DECLARATIONS

### Acknowledgements

We apologize to the authors that we could not cite due to space constraint. We thank Edith Heard, Maud Borensztein, Susana Chuva de Sousa Lopes, Hendrik Marks, Samuel Collombet and Bernhard Payer for discussions. We are grateful to the help of the KU Leuven FACS Core, Genomics Core and Mouse facility, Nucleomics Core, VIB/KU Leuven and SCIL.

### Availability of data and material

The accession number for RNA-seq data reported in this paper is GEO: GSE126229

### Authors’ contributions

A.J., I.T. and V.P. designed experiments; A.J., I.T., J.S., N.D.G., F.R., G.B., performed experiments; AJ., with input from I.T, S.K.T and F.R., analyzed sequencing data; Writing, A.J., I.T. and V.P. with input from all authors; Funding, V.P., J.C.M; Supervision, V.P.

### Ethics approval and consent to participate

Not applicable.

---

**Figure S1.**
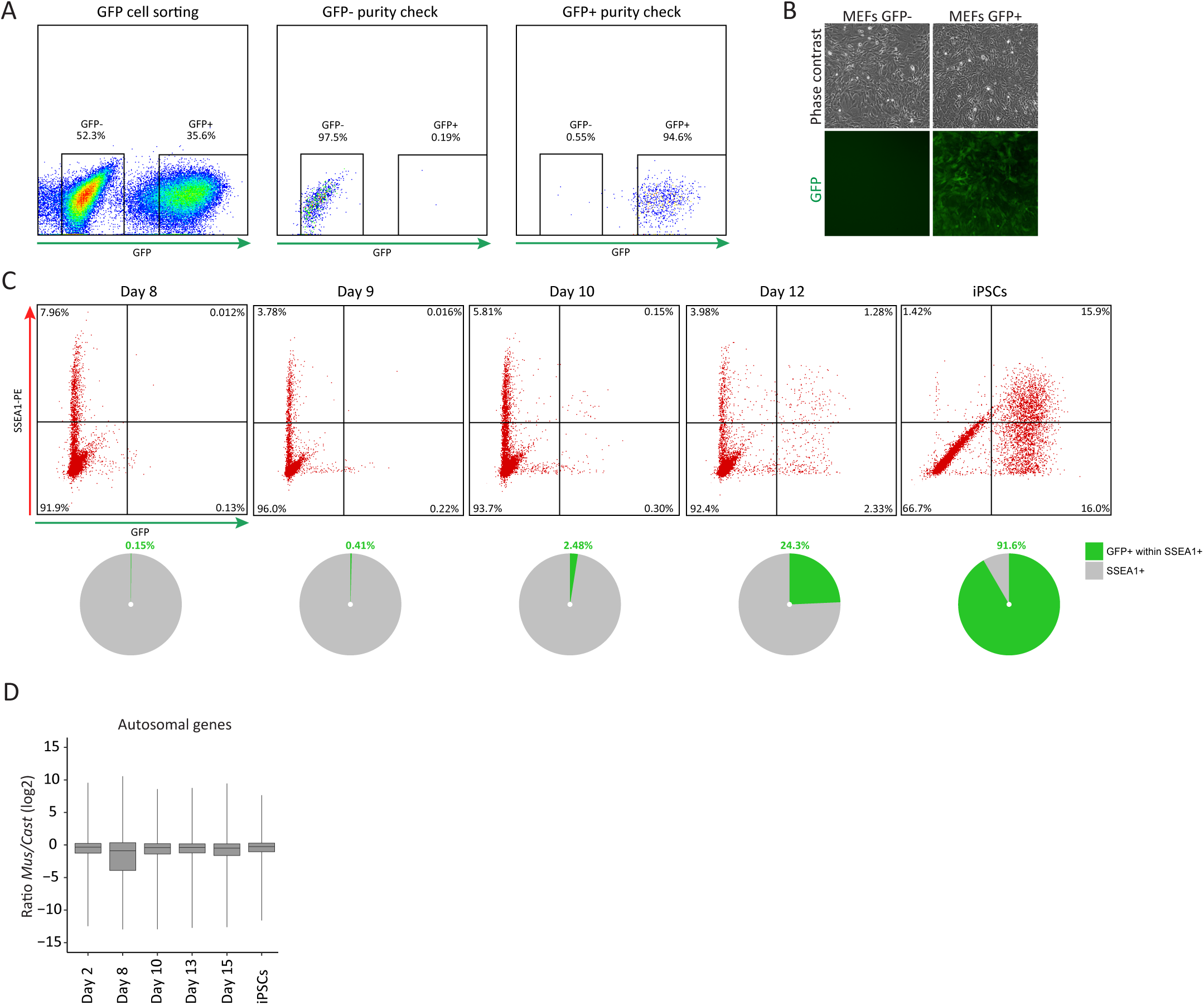
Analyses of *Mus*^Xi-GFP^/*Cast*^Xa^ reporter system. (A) Density plots representing the fluorescence-activated cell sorting for GFP- and GFP+ *Mus*/*Cast* MEFs and the purity check for both cell populations. (B) Phase contrast and fluorescent images of GFP*-* and GFP+ *Mus/Cast* MEFs populations. (C) Flow cytometry analysis of SSEA1+ and GFP+ cell population during reprogramming (day 8, 9, 10, 12 and iPSCs after four passages). The pie charts below refer to the proportion of GFP+ cells within SSEA1+ cell populations for each reprogramming time point. (D) Ratio of X-linked gene expression relative to autosomal genes (log2 transformed normalized counts) during reprogramming time points.

**Figure S2.**
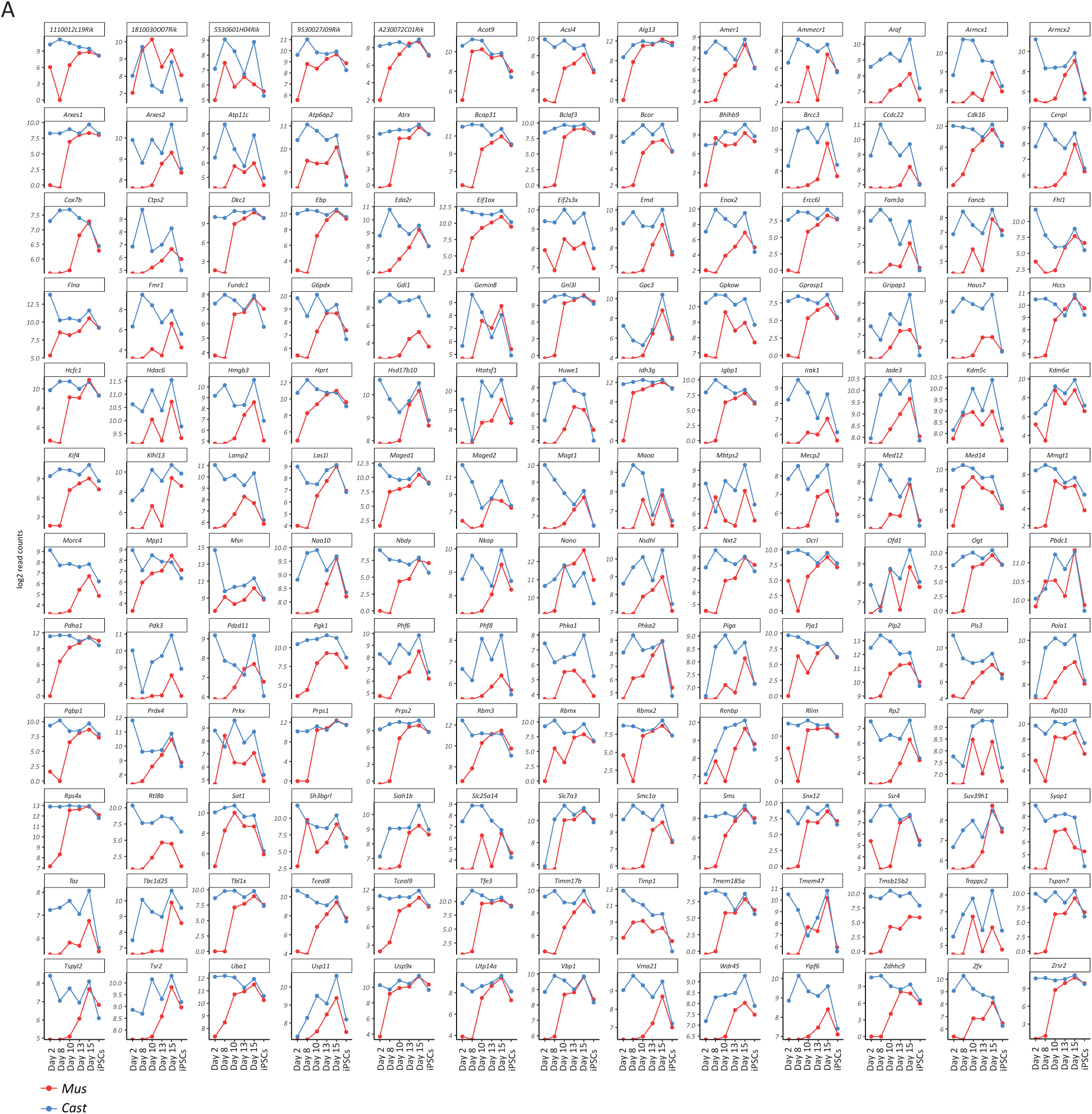
Reactivation kinetics of individual X-linked genes during factor-induced reprogramming to iPSCs. (A) Each graph represents the expression levels (log2 transformed read counts) of an informative X-linked gene listed in alphabetical order. The x-axis represents the different time points of reprogramming from day 2 to day 15 and iPSCs. Parental origin of the allelic expression is indicated in blue for *Cast* origin and in red for *Mus* origin.

**Figure S3.**
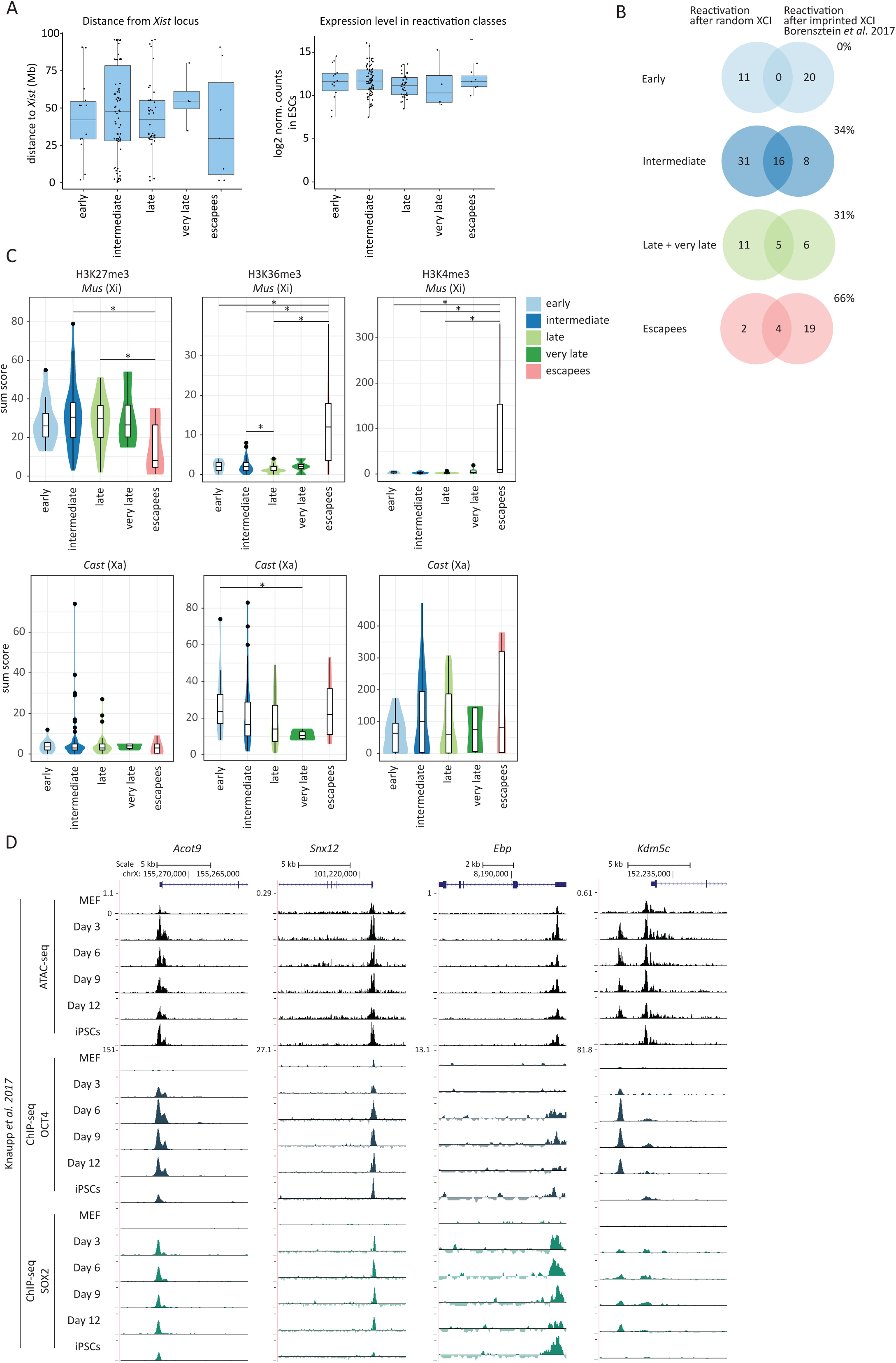
Possible predictors for XCR kinetics. (A) Distance to *Xist* TSS for each informative gene falling into each reactivation class (left) and for the gene expression level of each reactivation class (right) (log2 transformed normalized counts in ESCs). The *p*-values of a Kruskal-Wallis test comparing distance from *Xist* TSS to all sets of reactivation classes and gene expression level for each reactivation class were calculated and found to be not significant. (B) Comparison of XCR kinetics following iPSC reprogramming to that in the ICM. The number of overlapping genes for each category (early, intermediate, late + very late and escapee genes) in both data sets are showed inside the overlap region of the Venn diagrams. The proportion of genes overlapping in both datasets is shown. (C) Enrichment levels for H3K27me3, H3K36me3 and H3K4me3 occupancy for early, intermediate, late, very late reactivated and escapee genes in MEFs. The *p*-values of a Wilcoxon rank test comparing levels of enrichment between different gene reactivation classes are indicated with asterisks above the violin plots (****, *p*-value= 0.0001-0.001=extremely significant; ***, 0.001-0.01=very significant; **, *p*-value=0.01-0.05=significant; *, *p*-value≥0.05=not significant). (D) ATAC-seq, OCT4 and SOX2 ChIP-seq from SSEA1+ reprogramming intermediates (Knaupp *et al.* 2017) shown for *Acot9, Snx12, Ebp* and *Kdm5c*.

**Figure S4.**
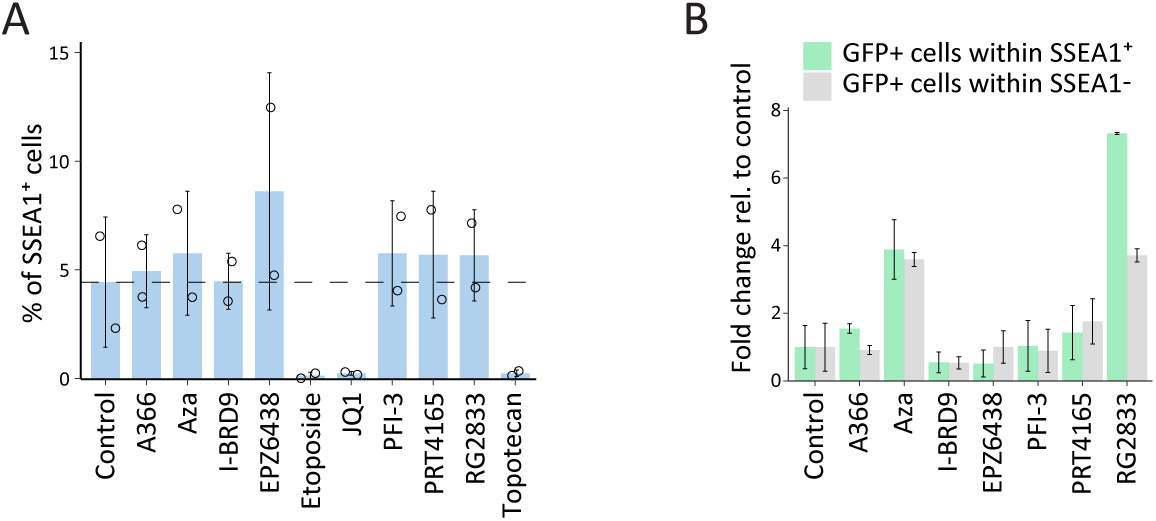
Epigenetic drug screening during reprogramming to pluripotency. (A) Proportion of SSEA1+ cells at day 10 of reprogramming for each individual drug inhibitor. (B) Fold change of GFP+ cells within SSEA1+ cell population relative to control (green) and fold change of GFP+ cells within SSEA1-cell population relative to control (grey) for each individual drug inhibitor.

**Figure S5.**
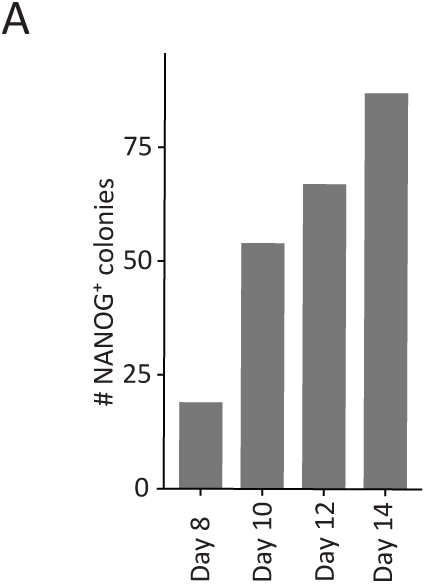
Number of NANOG+ colonies throughout reprogramming. (A) Number of NANOG+ colonies emerging during reprogramming to iPSCs. A colony is defined as four or more closely located cells.

